# Bayesian Reconstruction of Transmission within Outbreaks using Genomic Variants

**DOI:** 10.1101/213819

**Authors:** Nicola De Maio, Colin J Worby, Daniel J Wilson, Nicole Stoesser

**Affiliations:** Nuffield Department of Medicine, University of Oxford, Oxford, United Kingdom; Department of Ecology and Evolutionary Biology, Princeton University, Princeton, USA; Wellcome Trust Centre for Human Genetics, University of Oxford, Oxford, United Kingdom

## Abstract

Pathogen genome sequencing can reveal details of transmission histories and is a powerful tool in the fight against infectious disease. In particular, within-host pathogen genomic variants identified through heterozygous nucleotide base calls are a potential source of information to identify linked cases and infer direction and time of transmission. However, using such data effectively to model disease transmission presents a number of challenges, including differentiating genuine variants from those observed due to sequencing error, as well as the specification of a realistic model for within-host pathogen population dynamics.

Here we propose a new Bayesian approach to transmission inference, BadTrIP (BAyesian epiDemiological TRansmission Inference from Polymorphisms), that explicitly models evolution of pathogen populations in an outbreak, transmission (including transmission bottlenecks), and sequencing error. BadTrIP enables the inference of host-to-host transmission from pathogen sequencing data and epidemiological data. By assuming that genomic variants are unlinked, our method does not require the computationally intensive and unreliable reconstruction of individual haplotypes. Using simulations we show that BadTrIP is robust in most scenarios and can accurately infer transmission events by efficiently combining information from genetic and epidemiological sources; thanks to its realistic model of pathogen evolution and the inclusion of epidemiological data, BadTrIP is also more accurate than existing approaches. BadTrIP is distributed as an open source package (https://bitbucket.org/nicofmay/badtrip) for the phylogenetic software BEAST2.

We apply our method to reconstruct transmission history at the early stages of the 2014 Ebola outbreak, showcasing the power of within-host genomic variants to reconstruct transmission events.

**Author Summary:** We present a new tool to reconstruct transmission events within outbreaks. Our approach makes use of pathogen genetic information, notably genetic variants at low frequency within host that are usually discarded, and combines it with epidemiological information of host exposure to infection. This leads to accurate reconstruction of transmission even in cases where abundant within-host pathogen genetic variation and weak transmission bottlenecks (multiple pathogen units colonising a new host at transmission) would otherwise make inference difficult due to the transmission history differing from the pathogen evolution history inferred from pathogen isolets. Also, the use of within-host pathogen genomic variants increases the resolution of the reconstruction of the transmission tree even in scenarios with limited within-outbreak pathogen genetic diversity: within-host pathogen populations that appear identical at the level of consensus sequences can be discriminated using within-host variants. Our Bayesian approach provides a measure of the confidence in different possible transmission histories, and is published as open source software. We show with simulations and with an analysis of the beginning of the 2014 Ebola outbreak that our approach is applicable in many scenarios, improves our understanding of transmission dynamics, and will contribute to finding and limiting sources and routes of transmission, and therefore preventing the spread of infectious disease.

## Introduction

Understanding transmission is important for devising effective policies and measures that limit the spread of infectious diseases. In recent years, affordable whole genome sequencing has provided unprecedented detail on the relatedness of pathogen samples [1–4]. Consequently, accurately inferring transmission between hosts is becoming more feasible. However, this requires robust statistical approaches that make use of the full extent of genetic and epidemiological data available. Here, we present a new approach that makes use of within-host genetic variation and epidemiological data to infer transmission.

A number of approaches have been developed that reconstruct transmission from genetic data. The number of substitutions between samples from different hosts can be used to rule out transmission [5–7], or the phylogenetic tree of the pathogen samples can be used as a proxy for the transmission history [8, 9]. However, while the phylogenetic signal can be very informative of transmission, it can also be misleading [10, 11], due to within-host variation that can generate discrepancies between the phylogenetic and epidemiological relatedness of hosts, and can bias estimates of infection times [12, 13].

In recent years a number of methods have been proposed explicitly modelling both the transmission process and within-host pathogen genetic evolution to infer transmission events [11, 13–28]. Some of these methods use epidemiological data and genetic sequences from pathogen samples, and ignore within-host evolution and other causes of phylogenetic discordance with transmission history [14–19, 21–23]. Methods that explicitly model pathogen evolution within hosts and within an outbreak [13,20,24,25,27] generally assume, among other things, that samples provide individual and reliable pathogen haplotypes. This is often true for bacteria that are sampled and cultured before being sequenced, but it is mostly false for viruses and bacteria that are sequenced directly from samples without culturing. In fact, in these cases the sequencing process delivers reads coming from the different pathogen haplotypes that constitute the within-host pathogen population, and it is often very hard (if not impossible) to reconstruct complete haplotypes from these reads. In such cases, within-sample genetic variation is often neglected, and a single haplotype (which we call the consensus sequence of the sample) is built. While this procedure might lead to biases, it also certainly discards a very informative part of the available genetic data, because within-sample genetic variants can be very informative of epidemiological distance, direction of transmission, time from infection and transmission bottleneck intensity (see [29–32] and Figure 1). Furthermore, it is generally assumed that the pathogen does not recombine, so that a single phylogeny describes the evolutionary history of the whole genome, but this assumption does not fit highly recombinant pathogens such as HIV [33]. For these reasons, a few approaches have recently been proposed that use within-host genetic variants to reconstruct transmission [30, 32].

**Figure 1.**
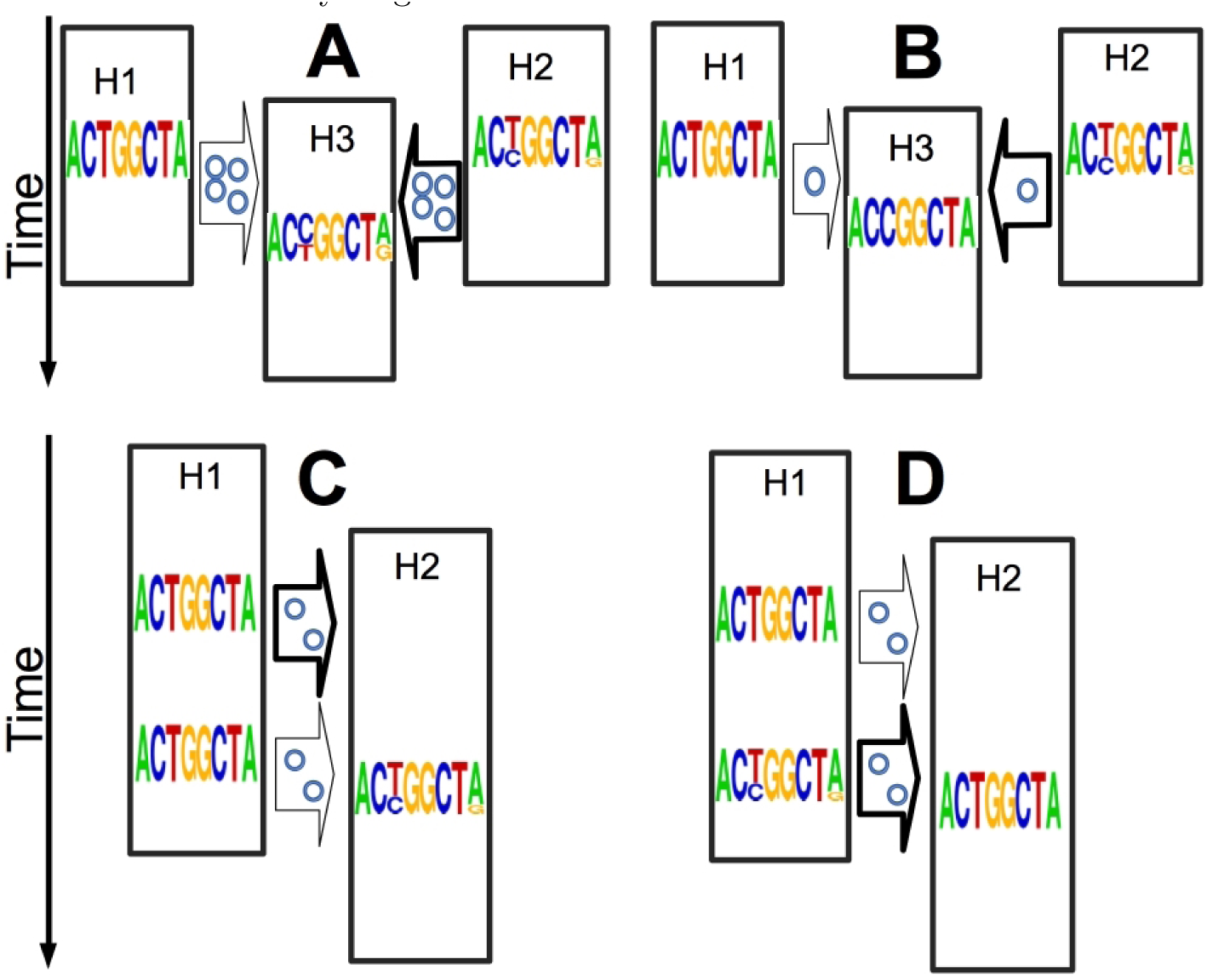
Examples of informativeness of within-host genetic variants. Here we show how within-host within-sample genetic variants can be useful without requiring pathogen haplotypes. Each string of letters (a frequency sequence logo [34, 35]) represents the collective genome of the pathogen at a certain point in time, as could be observed through deep sequencing. Multiple letters in the same column represent a genetic variant, with letter size representing allelic abundance. Time is on the Y axis, hosts are represented as black rectangles (a host is only active in the outbreak for the portion of vertical axis it occupies), and plausible transmission events as arrows. The posterior probability of different transmission events is represented by the arrow thickness. The number of little circles within arrows represents the inoculum size (transmission bottleneck). **A**) Shared genetic variants hint to epidemiological relatedness: the two top hosts (H1 and H2) are both possible infectors of the central host (H3), but H2 shares two genetic variants with H3, making it a likely infector of H3. Furthermore, the presence of shared genetic variants suggests a large transmission inoculum (a weak transmission bottleneck). **B**) A genetic variant of the same type of a substitution can hint to an infector: as before, but now H3 has a substitution (at third genome position, from T to C), which means that its within-host population is non-polymorphic at this position, but with a different nucleotide than the index case. This substitution is between the two nucleotides present at the same position in H2 (where this position is a genetic variant), consistent with H2 being the infector of H3. Also, this time the absence of shared genetic variants is indicative of a small transmission inoculum (a strong transmission bottleneck). **C-D**) The number of new genetic variants is informative of the age of an infection (but possibly also of the history of the pathogen population size within the host): in **C** the presence of non-shared variants in H2 suggests that the infection is older, while in **D** their absence suggests that the infection is younger.

Here, we propose a new Bayesian approach called BadTrIP (BAyesian epiDemiological TRansmission Inference from Polymorphisms) that not only uses within-sample genetic variants (from possibly multiple samples per host) to reconstruct transmission (including directionality and time of infection), but also combines this information with epidemiological data and an explicit model of within-host pathogen population evolution and transmission. We use the phylogenetic models with polymorphisms PoMo [36–38] to model population evolution along branches of the transmission tree; thanks to this, our transmission tree and phylogenetic tree are the same entity, and within-host evolution and recombination (resulting from a single primary infection, not multiple infections) do not create discrepancies that make statistical inference hard and computationally demanding [24, 25, 27]. We also explicitly model transmission bottlenecks, with one parameter defining the intensity of the bottleneck, and therefore the number of pathogen particles that establish a new population at transmission. Another feature of our approach is that, similarly to methods using within-host variants [30, 32], but differently from most other methods, we assume different genomic positions are unlinked; as such, our approach is expected to work well when recombination is strong enough to break linkage between genetic variants in the same host, or when very few high frequency variants and substitutions are expected per case, but could otherwise lead to poorly calibrated (excessively narrow) posterior probability distributions.

BadTrIP is implemented as an open-source package for the Bayesian phylogenetic software BEAST2 [39], and as such, it can be freely installed and used. We compare the performance of BadTrIP and the shared variants-based clustering (SVC) method of [30] on simulated data and on a real dataset from the early stages of the 2014 Ebola outbreak [40]. These applications show that BadTrIP has high accuracy to reconstruct transmission thanks to its explicit model of population evolution, the use of within-host genetic variants, and the inclusion of epidemiological data, and can provide important information to understand and limit the spread of infectious disease.

In the rest of the manuscript, we refer to a “host” as any entity that can contain and transmit a pathogen. Typically a host is a human within a community or nosocomial outbreak, or patients, but the concept of host can also be generalised for example to farms within a livestock outbreak. We will refer to the collection of all pathogens of the type under consideration within an individual host at a certain time as a “pathogen population” (for example all Ebola virions within an infected host, excluding non-Ebola pathogens and Ebola virions from other hosts). We will call a “pathogen unit” a single pathogen individual within a population, for example an individual bacterial cell or an individual virion. We call a pathogen population “polymorphic” at a particular genome position if pathogen units with different nucleotides at that position are present in the population; in this case, we also call the considered genome position a “genetic variant”.

## Results

### Modelling Within-Host Evolution, Transmission, and Sequencing

Methods to reconstruct transmission that account for within-host evolution usually have to deal with the complex task of modelling and inferring the discrepancies between the transmission tree and the pathogen phylogenetic trees [13,20,24,25,27]. We avoid this complication by adopting and adapting a substitution model, PoMo [36–38], that describes population evolution along the branches of a species (or population) tree. In this model, a virtual population, similar to a Moran model [41] without selection and with fixed population size, evolves by accumulating random changes in nucleotide frequencies (genetic drift, eventually resulting in the fixation of polymorphic sites), and new mutations resulting in new polymorphic sites. Different genome positions are modelled as completely unlinked.

The adoption of such a population genetic model within a transmission tree structure means that the phylogenetic tree and the transmission tree are now the same entity, and that each point of the tree represents the state of the pathogen population at a certain time within a host (Figure 2). Each bifurcation in the tree represents a transmission event, where the pathogen population splits in two groups: one remaining in the current host, and a small sub-population colonising a new host. We use a population bottleneck at time of transmission for the colonising branch to better model the transmission process.

**Figure 2.**
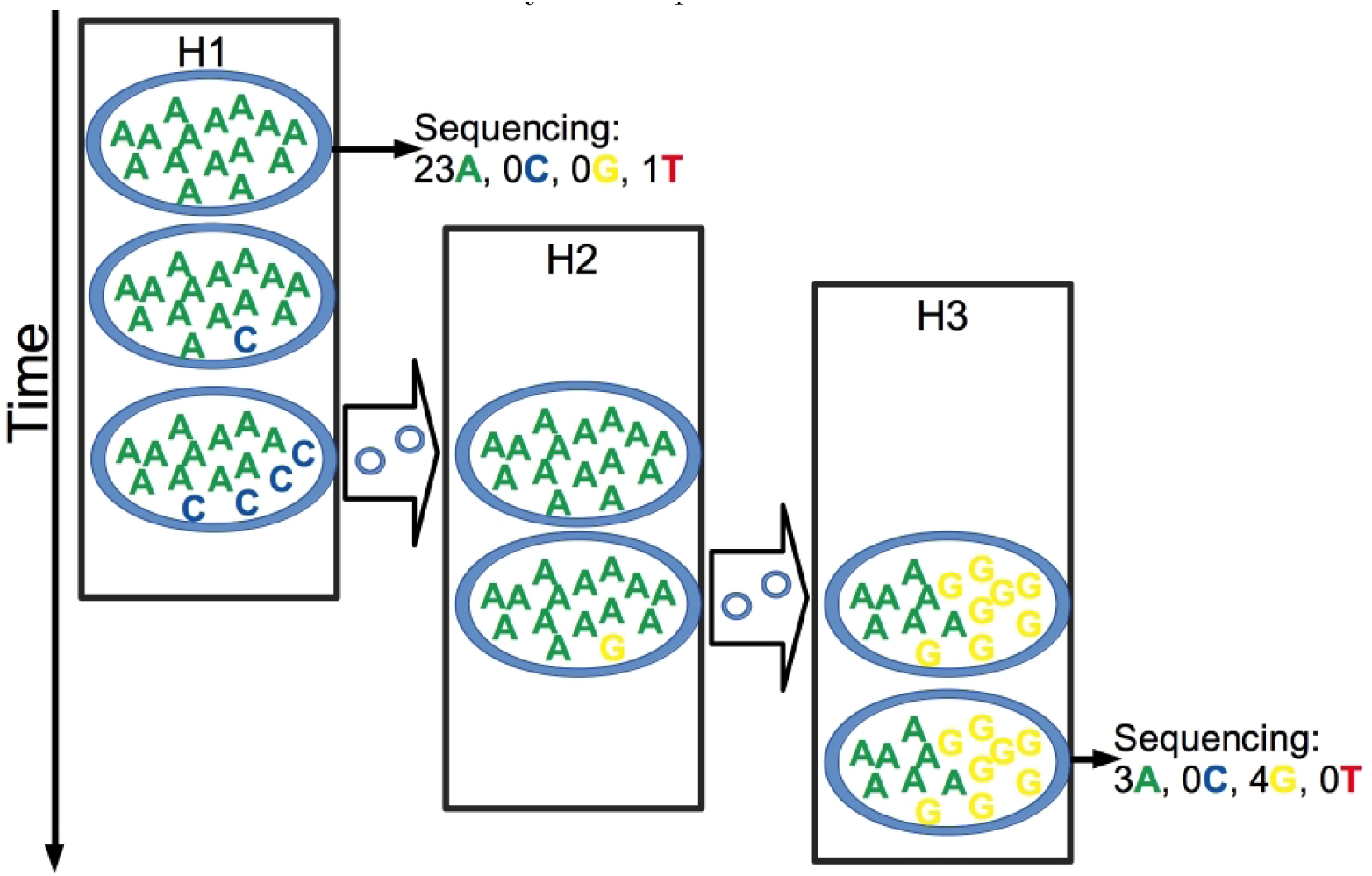
Graphical representation of the transmission, evolution and sequencing model. Here we describe some key aspects of our model. The figure depicts a possible evolutionary outcome for one position of the pathogen genome and the given transmission history. There are three hosts in this outbreak, represented by the black rectangles: H1 infects H2, which in turn infects H3. Time is on the vertical axis, and transmission events are represented by the thick arrows between hosts. Within each host, while it is colonised, the pathogen population consists of 15 units, each of which can have one of the four nucleotides at the considered position and at any time. For example, H1 starts off with all 15 pathogen units having an A, but during infection one of them mutates to C, and through genetic drift when H1 infects H2 it has 4 C’s and 11 A’s. While instantaneously only small changes can occur (one pathogen unit changing its nucleotide), along a time interval any number of changes can occur. As H2 is infected by H1, H2 is colonised by a copy of the pathogen population of H1, but the transmission bottleneck in this case causes one of the nucleotides to be lost, so that H2 is founded by a homogenous population of A’s. Within H2 again a mutation occurs and now a G is present in the pathogen population, but when H3 is colonised by H2 both nucleotides survive the transmission bottleneck, so H3 starts off with a polymorphic population. In the figure, H1 and H3 both have samples extracted and sequenced once, while H2 is not sampled at all. The sequencing process can result in any coverage (24 for H1 and 7 for H3 at the considered position). Furthermore, the observed nucleotide frequencies don’t necessarily exactly match the real nucleotides frequencies due to the randomness of read sampling, and because sequencing error can cause absent nucleotides to be observed at very low frequencies.

Our method uses two sources of information: epidemiological and genetic data. Epidemiological data is in the form of dates: the times when genetic samples are collected (it is possible to give any number of samples ≥ 0 for any host, even no sample at all) and a time interval for each host describing when it can contribute to the outbreak. Each host can only be infected, be sampled, and can infect other hosts within its time interval [13]. Genetic data from each sample is in the form of nucleotide counts: for each position of the genome, for a certain sample, the model expects the number of times each of the four nucleotides is observed in the reads (for example: 59 As, 0 Cs, 12 Gs, 1 Ts). We assume that reads are sampled with replacement from the pathogen population according to the (hidden) true nucleotide frequencies, and we model the sequencing error. This in particular means that sites without any sequencing coverage, or with very low coverage, are also allowed, and that differently from similar approaches (i.e. [30, 32]) we don’t required the specification of a minimum genetic variant frequency threshold.

While in our model we make the strong assumption that sites are completely unlinked, we test the performance of our approach with simulations in which we explicitly model within-host recombination events and we assume that a limited number of individuals in the pathogen population is sequenced. We even simulate scenarios in the total absence of recombination (complete linkage) to measure the robustness of our method. We simulate a broad range of scenarios: different transmission bottleneck severities (weak vs. strong), different amounts of genetic information, different recombination and mutation rates, different sequencing coverage levels, different sequencing error rates, and different virtual population sizes. We give further details on the model used and the simulations in the Materials and Methods section.

### Accuracy of Inference on Simulated Data

To test the accuracy of our new method BadTrIP in inferring transmission events, and to compare it to previous methods [30], we simulated pathogen evolution within outbreaks and sample sequencing, and we used different methods to reconstruct the transmission history from sequencing and epidemiological data. To simulate pathogen evolution, first we simulated an outbreak using SEEDY [42] (we used a fixed population of 15 hosts, one initial case, and an infection/recovery rate ratio of 1.43, see Materials and Methods); then, we translated the transmission history into a population history, and simulated within-population pathogen coalescent, recombination and mutation with fastsimcoal2 [43]. Throughout the simulations each host in the outbreak is sampled exactly once. We measure the accuracy of a method as the frequency with which the correct transmission source is inferred to be the most likely a posteriori. We also give a measure of the calibration of different methods by counting how often the correct source is in the 95% posterior credible set, defined as the minimum set of sources with cumulative probability ≥ 95% such that all sources in the set have higher posterior probability than all sources outside of it.

BadTrIP shows elevated accuracy in detecting the correct source of transmission (between 50% and 90%) and calibration (between 80% and 100%), in particular compared to the SVC approach (accuracy between 20% and 45% and calibration between 45% and 95%), see Figure 3. This shows that the use of epidemiological data and an explicit model of evolution can help to reconstruct transmission. In particular, comparing the base scenario with the one with almost no mutation, we see that BadTrIP accuracy drops from about 80% to about 50%, while the SVC accuracy drops from slightly more than 30% to about 20%; these drops approximately represent the contribution given by genetic data to the inference of transmission, while the difference between the two methods at almost no mutation (about 50% versus about 20%) shows the contribution of epidemiological information. Calibration of both methods increases as mutation rate decreases, one probable contributing factor being that as mutation rate decreases the effect of genetic linkage on the pathogen evolutionary dynamics decreases (neither method models genetic linkage). Similarly, the complete absence of recombination negatively affects calibration, but the difference is not dramatic (from about 90% to about 80% for BadTrIP, and even less for SVC) suggesting that even in the worst case scenario of complete absence of recombination BadTrIP can still provide meaningful inference and posterior distributions. Accuracy decreases with decreasing mutation rate, as is expected because of the reduced genetic information. However, increasing mutation rates to very high levels (to the point that about half the genome, of length 5kb, is polymorphic within the outbreak) does not seem to improve inference, probably because of saturation. Accuracy seems higher (around 10% difference) in the presence of a strong bottleneck (small inoculum) than a weak bottleneck (large inoculum), while calibration seems almost unaffected; this probably happens because, with strong bottlenecks polymorphisms are unlikely shared between hosts, and so polymorphisms leading to substitutions (see Figure 1B) become more informative for identifying infectors. An increase in coverage (from 40x to 100x) does not seem to bring improvement in accuracy or calibration; on the other hand, when a single uniform colony is sequenced (which is equivalent to reducing coverage to 1x, and therefore removing information on within-host genetic variation) seems to moderately reduce accuracy (≈ 10%) but not calibration. Introducing sequencing error (0.2% of mis-called bases, slightly more than what typical for high-throughput DNA sequencing [44]) accompanied by reduced coverage (20x) and genome length (1kb) still resulted in elevated accuracy (72.5%) and calibration (97.5%). Increasing the PoMo virtual population size (from 15 to 25, while the actual simulated population size remains 1000) had negligible effects on the inference.

**Figure 3.**
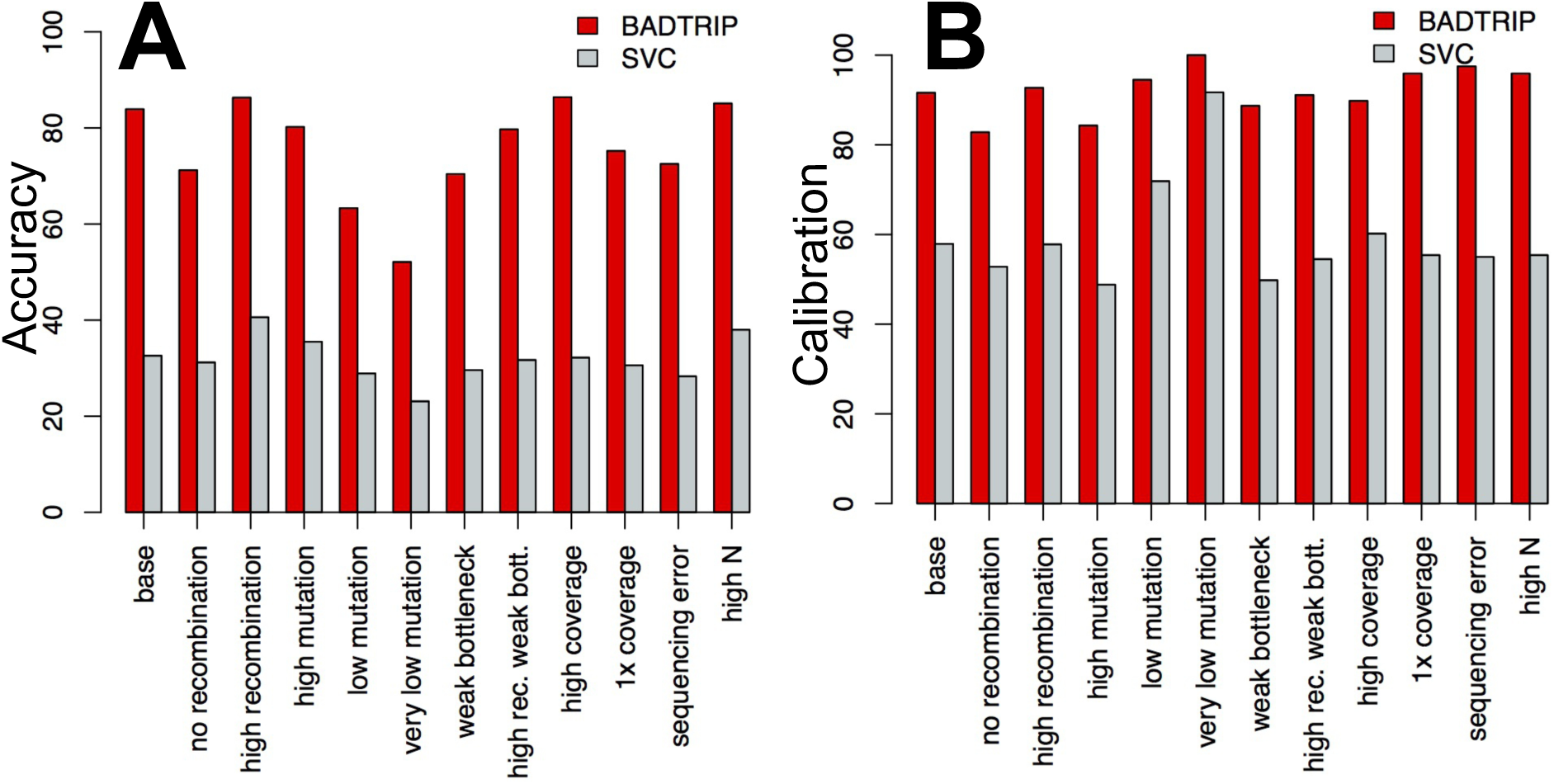
Accuracy and calibration of BadTrIP on simulated data. **A**) We represent accuracy as the frequency with which the correct simulated transmission event is more likely a posteriori than the alternatives. **B**) Calibration is the frequency with which the correct transmission event is in the 95% posterior credible set (the minimum set of sources with cumulative probability ≥ 95% such that all sources in the set have higher posterior probability than all sources outside of it). Bars represent percentages (from 0, worst, to 100, best) for BadTrIP (red) and the shared variants-based clustering (SVC) approach [30] (azure). On the x axis are different simulation scenarios with the first one, “base”, being the basic simulation scenario with 10-15 cases per outbreak, about 300-500 SNPs among all hosts, recombination 10 times stronger than mutation, complete bottleneck (no transmission of within-host genetic variants), read coverage of 40x, PoMo virtual population size of 15, actual pathogen population size of 1000, and genome size of 5 kb. All other scenarios are obtained from the base one changing one or two parameters: in “no recombination” the recombination rate is set to 0; in “high recombination” the recombination rate is 10 times higher; in “high mutation” the mutation rate is 10 times higher resulting in 2000-3000 SNPs per outbreak; in “low mutation” the mutation rate is 10 times lower resulting in 30-50 SNPs per outbreak; in “very low mutation” the mutation rate is 1000 times lower, resulting in 0-1 SNPs per outbreak; in “weak bottleneck” at transmission 5 pathogen units from the infector colonised the infected host, instead of just 1; in “high rec. weak bott.” both the recombination rate is 10 times higher and the founding population at transmission is made of 5 pathogen particles; in “high coverage” read coverage in sequencing is 100x instead of 40x; in “1x coverage” read coverage in sequencing is 1x instead of 40x; in “sequencing error” 0.2% of read bases are randomly modified to simulate sequencing error, coverage is reduced to 20x, and genome size is reduced to 1kb; in “high N” the PoMo virtual population size is 25 instead of 15.

BadTrIP also infers the time of infection. Calibration seems to increase with recombination, and to decrease with mutation (Figure 4), probably again an effect of our assumption of no linkage. Also, very high mutation rates seem to reduce the error in time inference, as do high coverage and virtual population size.

**Figure 4.**
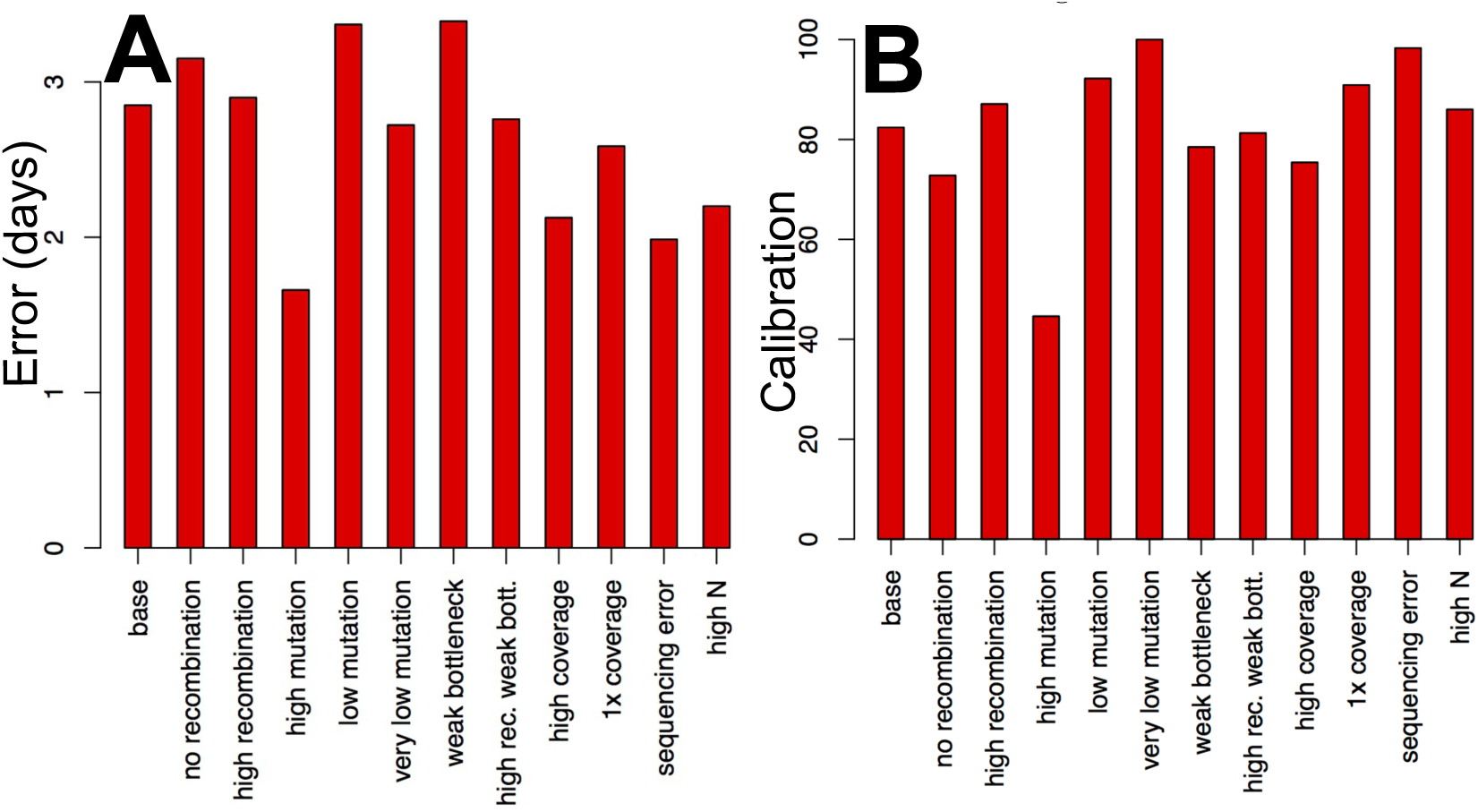
Error and calibration of BadTrIP inferring infection time from simulated data. **A**) Error (root mean square error) inferring times of infection with BadTrIP. The time unit is days, with a simulated transmission rate of 0.1 per day. **B**) Calibration (the percentage with which the true time of infection is within two standard deviations of the posterior median in the posterior distribution) of the inference of the time of infection with BadTrIP. Simulation scenarios are as in Figure 3.

The running time of BadTrIP is affected by the number of genetic variants present in the alignment and by the number of hosts present in the outbreak (Figure S1). The number of variants affect the number of likelihoods that need to be calculated at each MCMC step, while the number of hosts affects the size of the transmission/population tree (so both the computational and statistical complexity of BadTrIP). However, the time required to complete an analysis is not always a linear function of these two quantities: at low mutation rates BadTrIP requires similar times for different outbreak sizes. The reason is probably that with less data there is more uncertainty (in particular in the posterior distribution of the mutation rate), and so it takes longer to explore the the parameter space effectively. Overall, it takes a few hours to completely investigate an outbreak of moderate size (one or two dozen hosts) with BadTrIP.

### Analysis of the Early 2014 Ebola Outbreak in Sierra Leone

To demonstrate the applicability of BadTrIP and the advantage of using a model that combines epidemiological and within-sample genetic variation data, we use BadTrIP to infer transmission within the early cases of the 2014 Ebola outbreak in Sierra Leone. We use data published by Gire and colleagues [40] and previously analysed with the SVC method by Worby and colleagues [30]. One of the factors that make this dataset important to this study is the presence of within-host variants shared by multiple hosts, with one genetic variant that was even shared by eleven hosts [40]. While classical approaches based on consensus sequences would struggle to accommodate such data, in particular due to their assumption of strong transmission bottleneck that would not allow the transmission of variants, BadTrIP can accommodate such features, and such shared genomic variants are expected to increase the resolution of our transmission history inference. We investigate a collection of 62 samples with associated time and location of sampling. As observed by previous researchers, the number of substitutions (and more generally the number of SNPs) within this partial outbreak is very limited, and as such we expect to see a lot of uncertainty in the inference [30]; furthermore, all the samples were collected over a time interval of two months, and we assume transmission from a host to be possible from three weeks prior to three weeks following the sample collection, so the epidemiological data are also not very informative. Indeed, we see that most of the cases are inferred by BadTrIP to have a flat distribution of possible infectors, with highest per-infectee values generally under 30% posterior probability (Figure 5). However, we also see that BadTrIP identifies some pairs of infector-infectee with very high posterior probabilities (Figure S2). These pairs not only generally fit with the geographical epidemiological data, with most transmission with posterior probability *>* 50% happening within chiefdoms (with two exceptions discussed later), but also with the SVC inference [30]. Of these, transmission from EM119 to G3770 was inferred by Worby and colleagues [30] using consensus sequence genetic distance, while transmission from EM096 to G3679, from G3826 to G3827, from G3820 to G3838, from EM110 to G3809, and from G3729 to G3795 was inferred with the help of shared within-host genetic variants. All highly likely transmission pairs in [30] are also inferred by BadTrIP, but there are some highly likely transmission events inferred by BadTrIP that were not detected by SVC. For example, transmission from G3834 to G3817 is inferred by BadTrIP and is supported by a 3% frequency variant within G3834 that becomes fixed in G3817; however, such a variant fixation, attributable to the transmission dynamics described in Figure 1B, is not informative in the SVC method [30] and was further ignored due to the imposition of a 5% variant frequency threshold that we could avoid thanks to our explicit model of sequence evolution and sequencing error. Other cases similar to the latter are the inferred transmissions from EM110 to G3856, from EM110 to G3822, and from EM111 to G3724.

**Figure 5.**
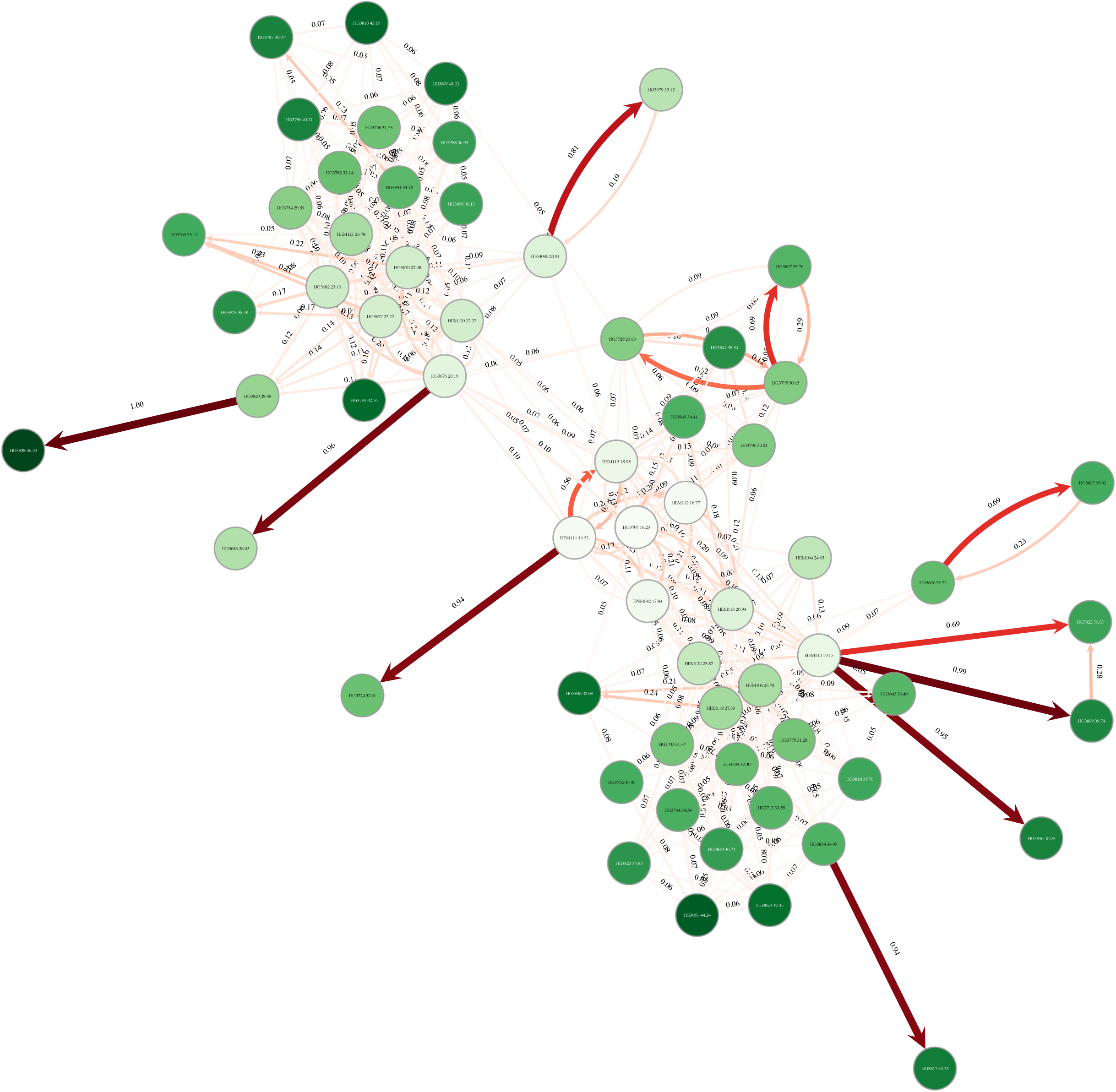
Inference of transmission in the early 2014 Ebola outbreak in Sierra Leone. **A**) Transmission events with posterior probability higher than 5% as inferred by BadTrIP. Circles represent hosts, while arrows are transmission events between hosts. The numbers next to arrows represent their posterior probability (between 0.0 and 1.0), as does their shade of red (from pale to dark red) and arrow thickness. Numbers within circles represent the inferred (posterior median) time of infection of the respective host, as also does the shade of green (from pale to dark green) of the circle. Time is expressed in days from the date of the first availability of the first host.

Cross-chiefdom transmissions inferred by BadTrIP with elevated posterior distributions are from EM110 in the chiefdom of Jawie, district of Kailahun, to G3856 in the chiefdom of Nongowa, district of Kenema; and from G3834 in the chiefdom of Kpeje to G3817 in the chiefdom of Jawie, both in the district of Kailahun. Neither of them had a high probability in [30], but they are both supported by low-frequency variants becoming fixed in the recipient.

Our inference of the sequencing error rate ɛ is extremely low (2 ⋅ 10^*−*7^ < *ε* < 7 ⋅ 10^*−*7^) consistent with the thorough filtering steps adopted by Gire and colleagues [40] prior to within-host variant calling.

### Discussion

Methods to infer transmission histories within outbreaks are important to determine the causes of transmission, to predict the most probable sources in the future, and therefore to inform policies preventing and limiting transmission. Genomic pathogen data from samples collected within an outbreak give the opportunity to observe at an unprecedented level of detail the genetic relatedness of pathogens from different cases. However, it is very hard to reconstruct the complete genome sequences of different pathogen units within the same host, even when sequencing output and read accuracy are elevated. The reason is that individual reads are generally shorter than the whole pathogen genome, and different reads usually come from different pathogen units. Within-host recombination, within-host mutation, and weak transmission bottleneck in fact cause the within-host pathogen population to be genetically varied. Within-host pathogen haplotype reconstruction is generally possible only when few high-frequency and diverged haplotypes are present, such as in the case of bacterial mixed infections with two diverged pathogen populations [45], or when within-host mutation is very high [46]. However, most methods to infer transmission from pathogen genetic data require full haplotypes, leading in many cases to loss of information (the within-sample genetic diversity) and consensus bias.

In recent years two methods have been proposed to infer transmission using not only genetic distances between consensus sequences, but also information about shared within-sample variants [30, 32]. In fact, within-host variants can be very informative regarding past transmission events (Figure 1), but it is not simple to accurately model pathogen population evolution within outbreaks and pathogen population sequencing. Here we present BadTrIP, a Bayesian approach to transmission inference that not only makes use of information regarding within-sample variants, but also implements an explicit model of pathogen population evolution within outbreak that allows inference of transmission direction and time, and allows the inclusion of epidemiological data that can further refine the inference of plausible transmission events. Compared to other similar methods [30, 32], our approach has the advantage of implementing an explicit model of pathogen population evolution and transmission, of allowing the inclusion of epidemiological data (sampling times and host exposure times), of not requiring minimum thresholds for within-host variant frequencies, of accounting for sequencing errors, and of being implemented as part of an open source phylogenetic package (BEAST2 [39]). Using simulations, we show that our approach achieves higher accuracy and calibration than SVC [30], and can reliably individuate likely transmission histories. Also, using a dataset of the early 2014 Ebola outbreak in Sierra Leone, and making use of information of within-sample variation and its evolution model, BadTrIP could individuate previously unidentified likely transmission events, including transmissions between geographic locations.

Despite these results, BadTrIP also has limitations, for example its model of genetic linkage. By assuming that all sites are unlinked, our model could be poorly calibrated in cases where there is no within-host recombination but high within-host mutation, causing strong correlations between inherited variants that are not expected in our model. However, we show in our simulations that our method is robust in a large variety of scenarios, including in the absence of recombination and with reads coming from few pathogen units. Another limitation is that our approach is generally not fast enough to deal with very large datasets, and, at the current stage, application is recommended to outbreaks with fewer than 100 cases. Also, BadTrIP is only applicable to the case where all hosts in the outbreak have been sampled, or at least observed. While this assumption is very common among transmission inference methods [11, 14–20, 23–26, 28] it also limits their applicability. Inferring possible non-sampled and non-observed intermediate hosts would probably lead to a significant increase in the statistical complexity and computational demand of BadTrIP (but see [13, 27]). Another scenario that is not accounted for in our model and should be therefore watched for is multiple infections of the same host (one host being infected by multiple sources, or from the same source multiple times). Another similarly looking and equally concerning problem is potential sample contamination. We recommend sequencing data to be searched for possible contaminations and multiple infections using methods such as PHYLOSCANNER [47] prior to be investigated with BadTRiP. We have also not accounted for selective pressure, which could sometimes introduce biases, for example creating homoplasies due to the same mutation appearing multiple times in different hosts, or the same polymorphism being maintained by balancing selection. However, our approach weighs information from both fixed substitutions and polymorphic variants, so the same mutation appearing in different genetic backgrounds will not be as nearly as biasing as for the SVC method. Furthermore, as our model is implemented in BEAST2, it is possible to specify a broad range of models of genomic variation in substitution rates which could at least partly account for the effects of selection. Finally, it is possible that errors in the bioinformatic processing of reads, for example mapping errors, cause the identification of the same spurious genetic variants in multiple hosts. We therefore encourage the investigations of genetic variants shard by many hosts to assess their biological plausibility. In the future we will work to solve some of the limitations of BadTrIP, in particular to reduce its computational demand and to model non-sampled non-observed hosts.

In conclusion, we have presented a new method that addresses the urgent need for software to efficiently and accurately analyse genomic and epidemiological data, in particular taking advantage of within-sample genetic variants to identify transmission pairs and reconstruct direction and time of infection. BadTrIP can be used in a broad range of outbreaks, and will be important for devising effective strategies to fight the spread of infectious disease.

### Software Availability

BadTrIP is distributed as an open source package for the Bayesian phylogenetic software BEAST2 [39]. It can be downloaded from https://bitbucket.org/nicofmay/badtrip/ or via the BEAUti interface [48] of BEAST2.

### Materials and Methods

### Model of Transmission

We model each host as a deme *d ∈ D* that can be colonised by a pathogen population, with total number of hosts-demes being *n*_*D*_. Each deme *d* is associated with an exposure interval limited by an introduction time *d_i_ ∈* (*−∞,* +*∞*] and a removal time *d_r_ ∈* [+*∞,+∞*), with *d*_*r*_ < *d*_*i*_ (we consider time backward as typical in coalescent theory); the host only contributes to the outbreak within this interval, which is determined by the epidemiological data. In the least informative scenario where no information on host *d* exposure is provided, it is assumed that *d* is exposed for the whole outbreak (*d*_*i*_ = + ∞ and *d*_*r*_ = − ∞). We will denote as ***E*** the collection of exposure times.

Each host-deme starts off as non-colonised and is colonised (infected) at some time *t*_*d*_ between *d*_*i*_ and the time that the first sample is collected from *d* (if no sample is collected from *d*, then we require only *t_d_ > d_r_*). Also, unless *d* is the first host to be infected in the outbreak, *d* is infected by another host in the outbreak *I*_*d*_ ≠ *d*, such that *I_dr_ < t_d_ < t_Id_*, that is, *d* is infected after *I*_*d*_ is infected, but before *I*_*d*_ reaches its removal time. If *d* is indeed the first case of the outbreak, then *I*_*d*_ is assigned the ∅ (we assume *∅ ∉ D*). We assume for simplicity that transmission between any pair of hosts and at any time is equally likely, as long as it is consistent with the epidemiological data.

Each host is also provided with a (possibly empty) set of samples, *S*_*d*_. Each sample *s* consists of a sampling time *t*_*s*_ and genetic data *G*_*s*_. Each sample *s* in *S*_*d*_ has to be collected after *d* is infected (*t_s_ < t_d_*) and before *d* is removed (*t_s_ > d_r_*). Assuming that the genome is *L* bases long, then the genetic data *G*_*s*_ of every sample *s* has to be in the form of a list of *L* quadruples, with for example the quadruple for genome position *i* being *G*_*si*_ = (*a_i_, c_i_, g_i_, t_i_*), the four positive natural values being the numbers of A’s, C’s, G’s and T’s observed at position *i* in the sample. If there is no read mapping to position *i* in sample *s*, then its quadruple is simply *G*_*si*_ = (0, 0, 0, 0). We denote the set of all sequencing data as *G*.

All hosts share a common parameter *B* (with real positive values) describing the intensity of the transmission bottlenecks associated with transmission events. Generally, the value of *B* can be inferred jointly with other model parameters, however its interpretation in terms of the size of the transmission inoculum is not straightforward. *T* denotes the transmission-population tree consisting of all sampling times, all infection times and all infectors of each host, and ***µ*** denotes the pathogen evolution model (described below).

We aim to sample from the following joint posterior distribution with a Monte Carlo Markov Chain approach:

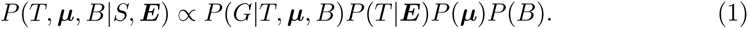

*P* (***µ***) and *P* (*B*) are the prior probabilities for respectively the substitution model and the bottleneck size, which can be chosen arbitrarily by the user. Instead, we ignore the prior for the transmission tree *P* (*T |**E***) as in [13]. *P* (*G |T, **µ**, B*) is the likelihood of the sequences given the genealogy and substitution model, and can be calculate as described below, using an adaptation of [36–38] to transmission trees. So once we calculate the likelihood *P* (*G T, **µ**, B*), we can use equation1 with an MCMC to infer a posterior distribution of infection times, infectors, bottleneck size and substitution model parameters.

### Model of Pathogen Evolution

Here, we make use of a phylogenetic model for population evolution, PoMo [36–38], to model mutation and drift in the within-host pathogen populations; also, we extend the model to include transmission bottlenecks and sequencing errors. Sequence evolution is usually modelled along phylogenetic trees, which can differ from the transmission tree [13]. However, PoMo describes evolution along species (or population) trees, and the population tree of a pathogen within an outbreak corresponds to the transmission tree *T* described in the previous section. If we consider the pathogen community within a host *d* as a population, we see that this population exists from time of infection *t*_*d*_, when it originates from a split with the population of its infector *I*_*d*_. So, transmission events corresponds to timed splits in the population tree, similar to the bifurcations of a species tree. However, one difference is that the split is asymmetrical, as we assume that the pathogen population size is not affected at *t*_*d*_ in *I*_*d*_, but at the start of the branch leading to *d* it undergoes a bottleneck of intensity *B*. All events in the tree are timed in real time (e.g., days) with some values fixed (for samples) and some values inferred in the MCMC (infection times).

We use a procedure very similar to the Felsenstein pruning algorithm [49] to calculate the likelihood of the genetic data over the tree. First of all, the substitution process along the branches of the transmission-population tree is not a simple DNA substitution process, but is similar to a 4-allelic Moran model [41] with mutation. We assume we have a continuous-time Markov process along each branch of the tree, where the state space is not made by the four nucleotides, as is typical, but by all 1- and 2-allelic states possible for a population of *N* units. Typical values of *N* that we use here are 15 or 25, that is, we describe evolution of a large population (possibly with billions of units) with a small virtual population of *N* units. Such an approximation generally lead to reasonably good results as long as we rescale the mutation rates between the real and the virtual population [36–38]. *N* here is not estimated, but is fixed by the user. Lower values of *N* are expected to reduce the computational demand of the method, but can result in lower accuracy. The states of our Markov process always include the four fixed states, where only one nucleotide is present in the population. In addition, they also include six groups of polymorphic states, where two nucleotides are present in the virtual population at the same site at the same time. Each group corresponds to one of the six unordered pairs of nucleotides (*{A, C}, {A, G}, {A, T}, {C, G}, {C, T}, {G, T}*) and contains *N −* 1 states: if the two nucleotides present in the population are *n*_1_ and *n*_2_, then such *N −* 1 states are the ones in which the population contains *i* times nucleotide *n*_1_ and *N − i* times nucleotide *n*_2_, for 0 *< i < N*. So in total our state space is of size 4 + 6(*N −* 1). Our substitution rate matrix is sparse, in that we only allow one unit in the virtual population to change at the time. So, from a fixed state with nucleotide *n*_1_, a instantaneous move is only possible to one of the three states with *N −* 1 times nucleotide *n*_1_ and one time any other nucleotide *n*_2_ different from *n*_1_. Such moves correspond to mutation events, and we represent their rates as *µ_n_*_1 *,n*2_. Instead, if we are already in a polymorphic state with *i* times nucleotide *n*_1_ and *N − i* times nucleotide *n*_2_, we only allow nucleotide counts to instantaneously change by one, so an instantaneous move is only possible to the state with *i* + 1 times nucleotide *n*_1_ and *N − i −* 1 times nucleotide *n*_2_, or to state *i −* 1 times nucleotide *n*_1_ and *N* + 1 *− i* times nucleotide *n*_2_ (one of these two latter states might be a fixed state). The instantaneous rate at which such changes happen is 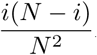 which corresponds to the rate of genetic drift (*R* represents the rate of drift in the virtual population in units of real time, which will depend on the generation time and is estimated by the model jointly with the other parameters). All other non-diagonal substitution rates are set to 0. All these states and rates constitute the substitution process ***E***.

The likelihood of *T* is calculated starting from the hosts in the outbreaks who don’t infect others (the leaves of the transmission tree). For such leaves, the likelihood is first calculated from the latest sample (if no sample is present, then the likelihood of such leaf at time of their transmission is 1 for every state). Given any state of our substitution process with nucleotides *n*_1_ and *n*_2_ with respectively abundances *i* and *N − i* in the virtual population (here for generality *i* can also be 0), given a sample and site at which the nucleotides with the highest coverage are *x*_1_ with coverage *c*_1_, and *x*_2_ with coverage *c*_2_ (we ignore the nucleotides with lower counts for numerical stability), then the likelihood of this state at this sample and site is approximated as:

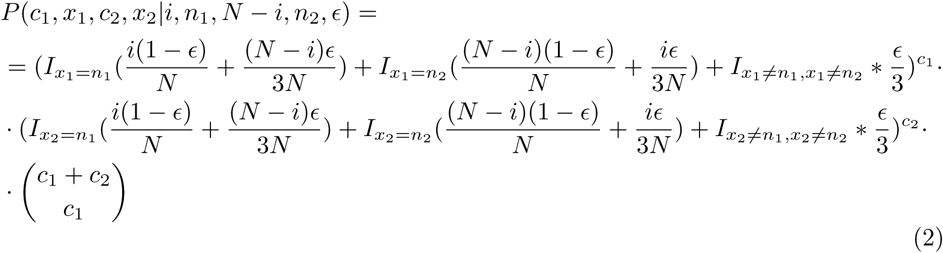

Where *ε* is a parameter describing the sequencing error rate. *ε* can be estimated with the other model parameters as we do with the real data and with the simulations including sequencing error. For all other simulations we set *ε* = 0. Along branches of *T*, the likelihood is updated using the matrix exponential of ***E***. At bifurcations (corresponding either to internal samples or transmission events) the likelihood is also updated according to the classical pruning algorithm, but at transmission events an extra step is added. A new drift-only substitution matrix ***E***_*D*_ is defined by setting the mutation rates in ***E*** to 0. Then, we describe a bottleneck as an branch of length *B* along which the population evolves under drift alone, that is, under ***E**_D_*. The length *B* does not count toward the branch lengths in real time, so that changing the intensity of the bottleneck does not affect the timing of the events in *T*. Under this model, a more intense bottleneck, corresponding to a small transmission inoculum, will be represented by a longer bottleneck branch, so a larger *B*. If we have a transmission event from *I*_*d*_ to *d* at time *t*_*d*_, we first calculate the likelihood within population *I*_*d*_ up to right before time *t*_*d*_ (likelihoods are updated backward in time), then within population *d* up to right before time *t*_*d*_, then we update the likelihood within *d* using the bottleneck branch, and finally we multiply the two likelihoods in *d* and *I*_*d*_ to obtain the likelihood in *I*_*d*_ right after *t*_*d*_ (again backward in time). This backward-in-time likelihood update process is terminated after the transmission event of the index case, and before its bottleneck we assume state equilibrium frequencies.

### MCMC operators

In addition to typical operator for *B*, *E* and ***E***, we also define five new operators for updating our transmission-population tree. The first operator modifies the transmission time *t*_*d*_ of a host *d*, without modifying any other parameter, not even the infector *I*_*d*_. The second operator picks a random non-index case *d* and, without modifying its infection time *t*_*d*_, picks a random new infector *I*_*d*_ among the ones compatible with infection time *t*_*d*_. The third operator is the same as the second, but first picks a new infection time for *d*, and then picks a new infector *I*_*d*_. The fourth operator exchanges a case *d* with its first infected case; the exchange swaps the infection times, and inverts the directionality of transmission between the two hosts. The last operator picks a random non0index case *d*, changes it infection time *t*_*d*_, and then picks a random new infector 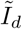 compatible with *t*_*d*_ and within the epidemiological neighborhood of *d* and *I*_*d*_ (for example *I*_*Id*_, other infectees of *I*_*d*_, or other infectees of *I*_*Id*_)

### Simulations of Pathogen Evolution

To test the accuracy of our new method BadTrIP in inferring transmission events, and to compare it with the SVC method [30], we simulated pathogen evolution within outbreaks and sample sequencing, and we used different methods to reconstruct the transmission history from sequencing and epidemiological data. To simulate pathogen evolution, first we simulated an outbreak using SEEDY [42] with a host population of 15 hosts and an infection rate of 0.1 per day, a recovery rate 0.07 per day, a conditionally accepting only outbreaks that achieve a minimum total of 10 infected case. Then, we translated the transmission history into a population history, assuming a within-host pathogen population size of 1000 and using fastsimcoal2 [43] to simulate pathogen coalescent, recombination and mutation with scenario-dependent parameters. Throughout all simulations each host was sampled exactly once.

We define a basic group of simulations (called “base”), and nine variants, in each of which one or two aspects of the base group of simulations is modified. In “base” we simulated about 300-500 SNPs (counting also variants present at very low frequency in just one host) or 45 substitutions per outbreak (which might be typical for HIV but high for many other pathogens), recombination rate 10 times higher than the mutation rate, complete bottlenecks (no transmission of within-host genetic variants), homogeneous read coverage of 40x, no sequencing error, PoMo virtual population size of 15, all equal mutation rates, and genome size of 5 kb. The nine variant settings are:

- **no recombination** - the recombination rate is set to 0.
- **high recombination** - the recombination rate is increased 10-fold.
- **high mutation** - the mutation rate is 10-fold higher resulting in 2000-3000 SNPs and about 385 substitutions per outbreak.
- **low mutation** - the mutation rate is 10-fold lower resulting in 30-50 SNPs and about 4-5 substituions per outbreak.
- **very low mutation** - the mutation rate is 1000-fold lower, resulting in 0-1 SNPs and 0 substitutions per outbreak.
- **weak bottleneck** - at transmission, 5 pathogen particles from the infector colonise the infected host, instead of just 1.
- **high recombination and weak bottleneck** - the recombination rate is 10-fold higher and the founding population at transmission is made of 5 pathogen particles.
- **high coverage** - read coverage is higher (100x instead of 40x).
- **1x coverage** - read coverage is extremely low (1x instead of 40x).
- **sequencing error** - read coverage is lower (20x instead of 40x), genome size is reduced (1kb instead of 5kb) and read bases are randomly modified to simulate sequencing error (0.2% of bases in reads are wrong).
- **high N** - the PoMo virtual population size is 25 instead of 15.

We ran 10 replicates for all scenarios, and 20 for “base”, “weak bottleneck” and “no recombination”. We ran the BadTrIP MCMC for 5 ⋅ 10^5^ steps for each replicate, sampled from the posterior every 100 steps and with a 20% burn-in. We specified in BadTrIP the true simulated sampling time and removal time of each host, while we specified as introduction time of each host its infection time minus one quarter of the mean duration of infection (so that the true infection time is contained within the exposure time of the host).

We measured accuracy as the frequency with which the correct transmission source is inferred by a method to be the most likely a posteriori. We also measured calibration as how often the correct transmission source is the the 95% posterior credible set (the minimum set of sources with cumulative probability ≥ 95% such that all sources in the set have higher posterior probability than all sources outside of it). In addition to performing inference from simulated data with BadTrIP, we also use the SVC method [30] which consists in selecting, for each host, the infector as the one with most shared variants, or, in the absence of shared variants, the one with the smallest consensus genetic distance. If multiple possible infectors score equally, they are assigned the same probability.

### The 2014 Sierra Leone Ebola Dataset

We use sequencing and epidemiological data published by Gire and colleagues [40] and analysed by Worby and colleagues [30]. In particular, we use information from sampling dates, nucleotide frequencies and sequencing coverage. We specify the introduction date (removal date) of each host as its sampling date minus (plus) 21 days. This means that we allow each host to be infected at most 21 days before it being sampled, and to infect others at most 21 days after being sampled. We ran the BadTrIP MCMC until an effective sample size of 1000 was reached for each parameter and for the posterior probability (requiring ≈ 3.5 million MCMC steps). to reduce the computational time required we subsampled the reads from each sample to obtain a per-base coverage of at most 100.

## Supporting Information

### S1 Text

**Supplementary Text S1.** The Supplementary text containing supplementary figures.

### S1 Data

**Supplementary Data.** Contains the xml script to replicate our Ebola analysis with BadTrIP, and a python script to replicate our simulations.

## Acknowledgments

D.J.W. is a Sir Henry Dale Fellow, jointly funded by the Wellcome Trust and the Royal Society (grant 101237/Z/13/Z). C.J.W. was supported by the Bill and Melinda Gates Foundation (grant OPP1091919). N.S. is currently funded through a Public Health England/University of Oxford Clinical Lectureship. N.D.M. was supported by the Oxford Martin School.

## Supplementary Text S1

**Figure S1.**
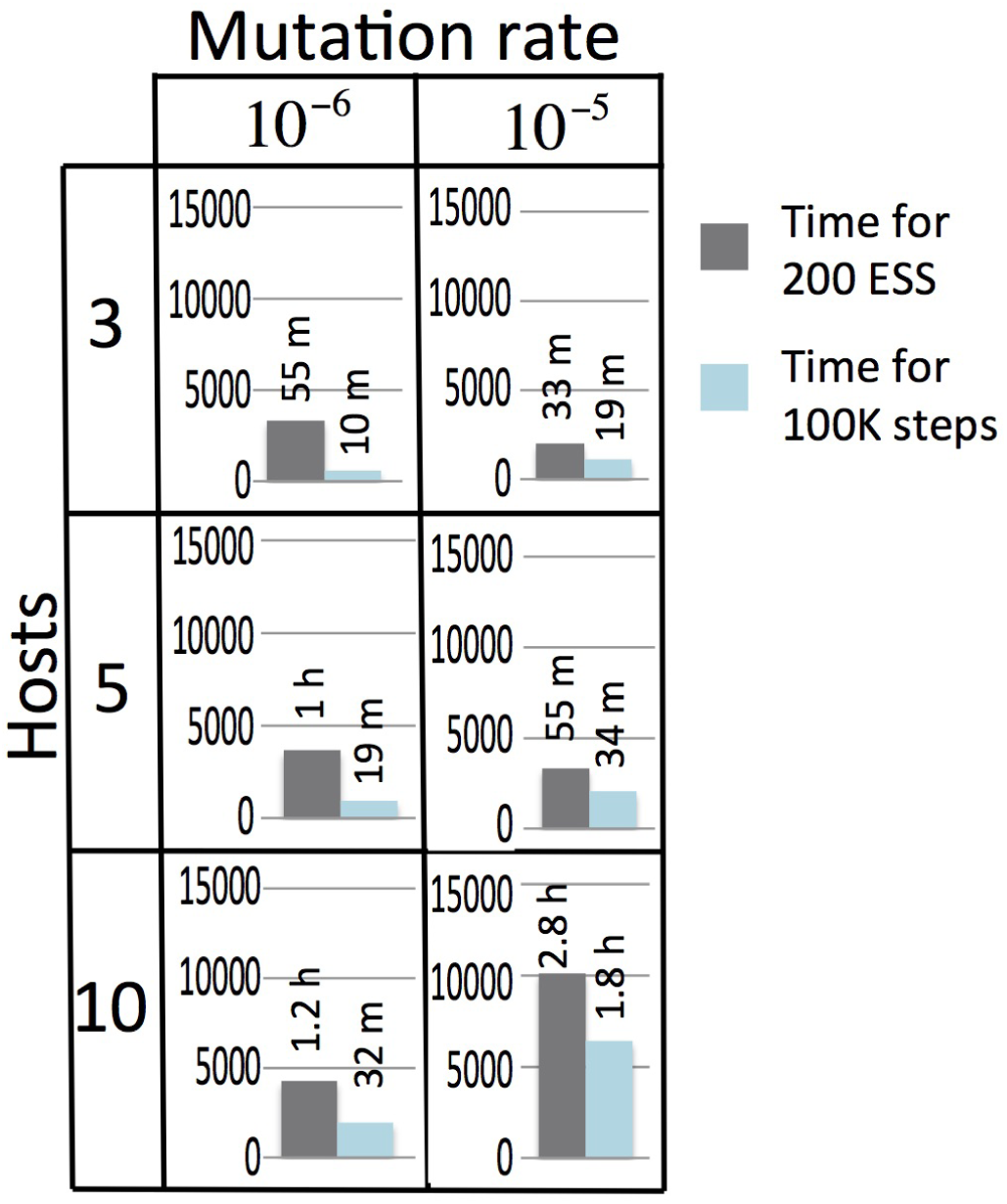
Computational demand of BadTrIP. Mean computational demand, in seconds, to run 10^5^ MCMC steps (blue) and to achieve an effective sample size of 200 for the posterior probability (grey) in BadTrIP. Each barplot represents the mean over 10 simulations. The three rows in the table represent the number of hosts in the simulated outbreaks (3, 5 or 10) and the two columns represent different mutation rates (10^*−*5^ corresponds to the “base” scenario in Figure 3 while 10^*−*6^ corresponds to the “low mutation” scenario in Figure 3). On top of each bar is the same running time but represented in minutes (“m”) or hours (“h”).

**Figure S2.**
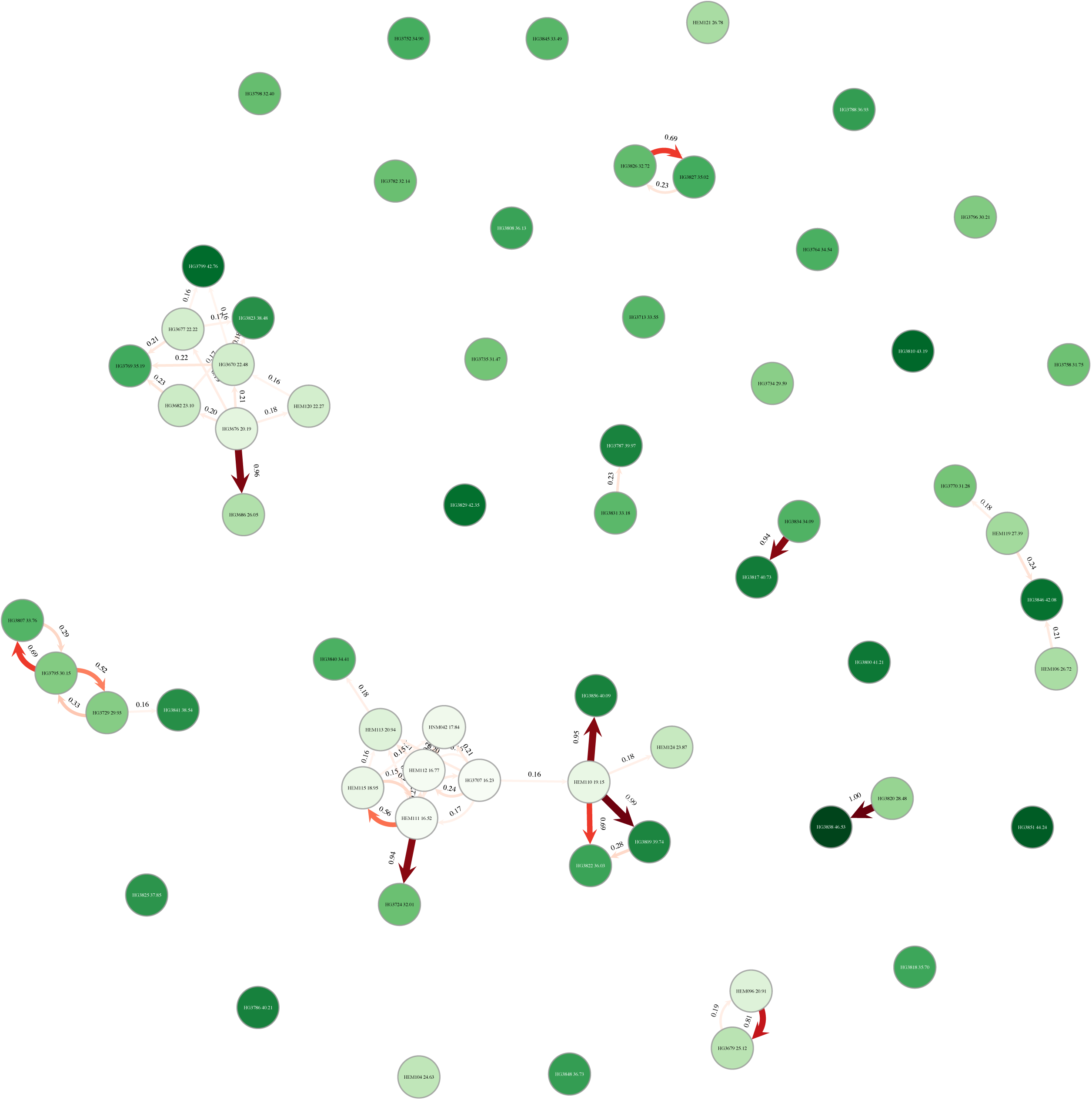
Inference of transmission in the early 2014 Ebola outbreak in Sierra Leone, only high-probability transmissions. Transmission events with posterior probability higher than 15% as inferred by BadTrIP. The notation is as in Figure 5

